# Yeast PAF1 complex restricts the accumulation of RNA polymerase III and counters the replication stress on the transcribed genes

**DOI:** 10.1101/339499

**Authors:** Pratibha Bhalla, Dipti Vernekar, Ashutosh Shukla, Benoit Gilquin, Yohann Couté, Purnima Bhargava

## Abstract

Many regulatory proteins and complexes influence transcription by RNA polymerase (pol) II. In comparison, only a few regulatory proteins are known for pol III, which transcribes mostly house-keeping and non-coding genes. Yet, pol III transcription is precisely regulated under various stress conditions like starvation. We used pol III transcription complex components TFIIIC (Tfc6), pol III (Rpc128) and TFIIIB (Brf1) as baits to identify potential interactors through mass spectrometry-based proteomics. A large interactome constituting known chromatin modifiers, factors and regulators of transcription by pol I and pol II revealed the possibility of a large number of signaling cues for pol III transcription against adverse conditions. We found one of the pol II-associated factors, Paf1 complex (PAF1C) interacts with the three baits. Its occupancy on the pol III-transcribed genes is low and not correlated with pol III occupancy. Paf1 deletion leads to higher occupancy of pol III, γ-H2A and DNA pol2 but no change in nucleosome positions. Genotoxins exposure causes pol III but not Paf1 loss from the genes. PAF1C promotes the pol III pausing and restricts its accumulation on the genes, which reduces the replication stress caused by the pol III barrier and transcription-replication conflict on these highly transcribed genes.

## Introduction

Eukaryotic transcription is accomplished by three multi-subunit RNA polymerases (pol)s; pol I, pol II and pol III; of which pol III has seventeen, the largest number of subunits. but the simplest transcription complex. It consists of pol III, the basal factors TFIIIC, TFIIIB and TFIIIA-the 5S-specific assembly factor (1, 2). In yeast, the six-subunit assembly factor, TFIIIC binds the intra-genic promoter elements and recruits the three-subunit initiation factor IIIB at the −30 bp position, upstream of the target genes, while the 17-subunit pol III is recruited by the initiation factor TFIIIB (3). Pol III transcription units are small, giving nascent transcripts of generally 100 bases average length (4). High transcription activity in vivo (5) is proposed to interfere with replication fork progression on these slow-replicating genes. The resultant fork pausing at the pol III-transcribed genes makes them prone to the replication stress-induced DNA damage (6, 7).

It has been known since long that DNA-protein and protein-protein interactions are instrumental in the gene regulation process. Extensive research on mechanisms regulating the transcription by pol II has helped understand the details of mRNA production in the eukaryotic cell. In contrast, so far only a few regulatory proteins are known for pol III transcription. Yet, pol III transcription is repressed under a variety of stress conditions including nutrient starvation (8–10). A number of accessory factors, which do not show sequence-specific DNA-binding have been proposed to influence the pol III transcription, without apparent gene specificity (11). Maf1, p53, RB, Sub1, p300, the La protein, NF1, Nhp6, TFIIS and TFIIIE (10–15) are few such examples. In the absence of known upstream regulatory elements (16) or classical, DNA-binding activators/repressors for the pol III-transcribed genes in yeast, chromatin related epigenetic mechanisms have been proposed and shown to be their regulators (17). With binding of TFIIIB in the upstream region, TFIIIC not leaving the gene body during transcription or under repression (18, 19) and the pol III possibly recycling at high rate on the genes (20, 21), the whole TC components remain in close proximity of each other all throughout the transcription process (22, 23). Therefore, proteins interacting with any of the components in one part of the complex may influence the activity and topology of the whole complex in space.

Proteomic approaches have been used as strong tools to find the global interacting partners of many complexes (24–27). Therefore, in order to find the possible regulatory partners of the pol III TC, we followed AP-MS proteomic approaches (28) wherein we selected one subunit of each component to pull down the interacting partners in its transcriptionally active and repressed (nutrient starvation) states. A large number of proteins, including subunits of several known functional complexes, were found associated with the pol III TC. Interestingly, among the various machineries co-immunoprecipitating with the pol III TC, we identified a known transcription factor of pol II, the Paf1 complex (29), which showed gene-specific, low levels on the pol III-transcribed genes. Paf1 complex associates with pol II (30) and has two main activities on pol II-transcribed genes (31, 32). It is required for transcription elongation by pol II and pol I (33–35) and for coordinating the co-transcriptional histone modifications on the gene body nucleosomes (30, 36). However, its role in proper 3’-end formation of the transcripts (37) and several other non-transcriptional processess like DNA repair, cell cycle, gene silencing etc. (31, 38) are not well understood yet.

Phosphorylation of the Ser139 of mammalian histone H2A.X (and Ser129 residue of yeast H2A) generates γ-H2A.X (yeast γ-H2A), the marker of the DNA damage (39, 40). We found low levels of γ-H2A and the DNA pol2 subunit of the pol ε on the yeast pol III-transcribed genes, which increased upon exposure to the genotoxin HU and upon Paf1 deletion. In the *paf1*Δ cells, pol III levels increased several fold on all the tested genes but only a small increase in tRNA levels were seen. Paf1 influences the effect of the genotoxin HU but not MMS and restricts pol III occupancy levels on the genes. These results along with low levels of active state H3K4 and H3K36 histone methylation marks found near pol III-transcribed genes suggest a repressive influence of Paf1 on pol III-transcribed genes, which may serve to protect these highly transcribed, known replication fork pause sites (6, 7) against the DNA damage.

## Materials and Methods

### Yeast strains, plasmids, media and growth conditions

Yeast strains and primers used in this study are listed in the Table S1 and Table S2 respectively. Cells were grown normally in YEP (yeast extract and peptone) medium containing 2% glucose at 30^0^C. Cells were nutritionally deprived by shifting to 0.15X YEP without any carbon source after attaining the OD_600_ of 0.8 and allowed to grow for 2 more hrs at 30^0^C, before harvesting. The genotoxin effect was followed by exposing the mid log phase cells to final concentration of 0.05% (v/v) MMS or 200mM HU for 2 hours, before harvesting. Brf1 was FLAG-tagged according to Gelbart et al, 2001 (41) and other taggings were performed by chromosomal integration of PCR amplified cassettes as described in Janke et al 2004 (42).

### Antibodies

Anti-FLAG antibody (F7425) and anti-FLAG M2 agarose beads (A2220) were purchased from Sigma. Anti-Myc tag and anti-HA tag antibodies used for for ChIP and IP were from Millipore (05-724 and 05-904 respectively). Anti-myc, anti-HA and anti-FLAG antibodies used in western probings were from Santa Cruz ((9E10) SC-40 and (Y-11) SC-805) and Millipore (MAB3118) respectively. IgG sepharose beads were from GE Healthcare. Anti-H3 antibody (abcam 1791) and antibody against phosphorylated H2A S129 (ab15083) were from Abcam.

### Sample preparation for proteomics experiments

The transcription complex of pol III is a compact assembly, which covers the complete gene body of the short transcription units (18, 22, 23, Fig. S1A). We used one subunit each of pol III (Rpc128), TFIIIB (Brf1) and TFIIIC (Tfc6) as bait in our proteomic experiments. Yeast cells individually expressing FLAG-tagged baits were grown at 30^0^C in YEPD media. Cells grown under the nutrient-enriched or –deprived conditions were harvested and lysed mechanically by vortexing with glass beads in lysis buffer (50 mM HEPES [pH 7.5], 150 mM NaCl, 2mM MgCl_2_, 0.5mM EDTA, 0.1% NP-40, 10% glycerol) at 4^0^C. Lysates were centrifuged at 10000xg for 10 min and supernatant was treated with DNaseI before incubating with IgG sepharose beads at 4^0^C for 2 hrs for pre-clearing. Cleared lysates were incubated with anti-FLAG M2 agarose beads for immunoprecipitation (IP) at 4^0^C for 3 hrs. Pol III control samples were prepared by incubating the lysates with IgG sepharose beads for mock whereas for Brf1 and Tfc6 control samples, lysates of the untagged yeast strains of the same genotype were incubated with anti-FLAG M2 agarose beads for immunoprecipitation (IP) at 4^0^C for 3 hrs. Beads were washed with lysis buffer and affinity purified proteins were released from the beads by FLAG peptide elution. The 1/10^th^ volume of eluate was resolved on 10% SDS-PAGE and visualized by silver staining for checking IP. A typical profile of IP samples used for analysis is shown in the Fig. S1B-D. Eluates were boiled, loaded on 10% SDS-PAGE and allowed to stack by stopping the run when tracking dye entered 0.5 cm in resolving gel (Fig. S1E). Gel was stained with coomassie brilliant blue R-250 and gel area containing protein was cut into 1mm^3^ pieces to process for Trypsin digestion.

### Mass-spectrometric analysis

Resulting peptides were analysed by nano LC-MS/MS. Further details of proteomic experiments, and processing of Data output files, generation of identified protein lists and bioinformatic analyses of positive hits are elaborated under Supplementary methods in the accompanying File S1.

Differential analysis of control and positive samples was performed using extracted specific spectral counts (SSC). For potent stringency, SSC of negative control were calculated as the sum of three biological replicates whereas for SSC of individual baits, two biological replicates were summed. To be considered as a pol III TC binding partner, a protein was to be identified with a minimum of 4 summed SSC and only in a bait IP (not in control) or enriched at least 5 times in the sample as compared to the control one.

### Immunoprecipitations (IPs) and Western analysis

Cells were grown to logarithmic phase YPD at 30^0^C and harvested, washed with ultrapure water and lysis buffer (50 mM HEPES [pH 7.5], 150 mM NaCl, 2mM MgCl2, 0.5mM EDTA, 0.1% NP40, 10% glycerol). The lysis buffer was supplemented with 1mM PMSF, 2mM sodium orthovanadate, 1mM sodium fluoride and 1x protease inhibitor cocktail (proteCEASE G-Biosciences) during lysis by vortexing with glass beads in mini bead beater (Biospec) at 4^0^C. Immunoprecipitations were performed with anti-FLAG M2 agarose beads or anti-HA antibody (Millipore 05-904). Affinity purified proteins were released by boiling in Laemmli buffer and resolved on SDS-PAGE. Proteins were transferred to PVDF membrane and analysed by Western probing. Unless otherwise stated, all the co-IPs were performed with pairing of Flag and HA tags. The blots were sequentially probed with or without mild stripping after first probing and one antibody at a time. IP and co-IP bands are marked for each pull-down, shown in representative Western images.

### Chromatin Immunoprecipitation (ChIP) and Real Time PCR

ChIP and Real Time PCR estimations were performed as described earlier (43). Cells were grown in YPD at 30^0^C and fixed with 1% formaldehyde. Cells were washed thoroughly and lysed mechanically by glass beads in lysis buffer (50 MM K-HEPES pH 7.6, 1% Triton-X 100. 0.1% sodium deoxycholate, 2mM EDTA, 150 mM NaCl) at 4^0^C. Chromatin was sheared to average size of 300 bp by sonication and used for immunoprecipitation with corresponding antibodies (anti-Myc, anti-HA from Millipore) along with Protein A or Protein G agarose beads (GE Healthcare). Beads were washed in wash buffers and eluted at 65^0^C in elution buffer (10 mM Tris-Cl pH 8, 2mM EDTA, 200 mM EDTA, 1% SDS) for 30 min. The ChIP DNA was precipitated by ethanol after deproteinization with phenol, dissolved in TE and quantified in 10 μl reaction mixtures by a real-time PCR method based on SYBR green chemistry using Roche LC480 platform. The subtelomeric region of the right arm of the chromosome VI (TEL VIR) was used as a control region. Occupancies were calculated as the relative enrichment above the control values and normalized against the value for mock immunoprecipitation.

The “Fold enrichment method” was used to calculate the relative enrichments of DNA pol 2 over the background using TelVIR as internal control region. In this method, the Ct values obtained from the ChIP and mock samples were first normalized with Ct values obtained from TelVIR region and then by Ct values of Input (88). Average and scatter from minimum three independent estimations are plotted. Genes showing non-significant changes with p value >0.05 are marked with a dot wherever applicable. Rest of the genes have significant p values (<0.05).

### Northern blotting for tRNA estimations

The method used by Karkusiewicz et al, 2011 (44) was used for tRNA estimations by Northern blotting. Per lane 10 μg of total RNA was resolved on 15 % Urea-PAGE and transferred from gel to Hybond N+ membrane (GE Healthcare) by electroblotting in 1x TBE, cross-linked by UV radiation (1200 mJ/cm2). The membrane was pre-hybridized in prehyb/hyb buffer (2x SSC, 1X Denhardt’s solution, 0.2%SDS) and hybridized at 37^0^C in the same solution with oligonucleotide probes (listed in the Table S2) labeled with [γ^32^P]-ATP by T4 polynucleotide kinase (New England Biolabs). The membrane was washed 3 times for 15 min with wash buffer (2X SSC and 0.2% SDS) at 42^0^C and exposed to phosphorimaging screen (Fujifilm). Bands were visualized in phosphoimager (BIORAD) and intensities were quantified using Image Gauge software from Fuji.

### Genome-wide data analysis

The processed tab files containing forward and reverse tag counts, with their respective chromosomal location were used for the analysis of the ChIP-exo data (45) on the occupancy of the PAF1C subunits on the tRNA genes were downloaded from GSE83348. The open source “Bio toolbox” from Timothy J. Parnell was used to map binding sites of all the subunits of Paf1 complex around the TSS of the tRNAs genes. The genome-wide occupancy of Rtf1 on the tRNA genes in the yeast genome was analysed also from the published ChIP-seq data (46). The reads uniquely aligned to sacCer3 yeast assembly with mapping quality scores >30 were processed for further analyses. The “bamCompare” function from deepTools package was used for Library size adjustments (SES method) and log_2_ ratio of IP vs Input was calculated (47). The “plotProfile” and “plotHeatmap” functions of deepTools were used for plotting the average occupancy profiles and heatmaps (47).

All Raw data files for nucleosomal DNA (48) from the wild type and *paf1*Δ cells as treatment group and whole genome fragmented DNA as a control group were downloaded from NCBI GEO under accession no. GSE44879. The two sample analysis was performed on the nucleosomal occupancy data by employing Affymetrix Tiling Analysis Software (TAS) v1.1. Data were quantile normalized in pairs using only perfect match probes with a bandwidth of 30. Resulting bar files were used for Interval analysis with p-value threshold of 0.05, minimum run of 50 and maximum gap of 20 probes. Processed MNase-seq data files on histone methylations were downloaded under accession no GSE61888 from NCBI GEO. The plotHeatmap” function from deepTools was further used for mapping of respective histone methylations (49) to yeast tRNA genes..

### Statistical analyses

Each measurement was repeated several times and average with scatter for minimum three biological replicates was taken for plotting the graphs. Statistical significance of a difference between the two values was calculated using two-tailed unpaired Student’s t-test. The insignificant changes (p>0.05) if any, in every set of data are marked in every Figure panel. Rest of the comparisons are significant (p<0.05).

## Results

### Many interactors of pol III TC are common between active and repressed states

Using three components of the pol III TC as baits in our proteomic experiments, we found a large number of potential binding partners of pol III TC. A total of 411 proteins in the active state and 405 proteins in the repressed state could be taken as positive hits, a majority of them being common between the two states (Fig. 1A). A large number and variety of associated protein complexes were detected in this study, which include in the active state, 30 out of total 44 interactors of Rpc160, reported previously (50). Venn intersections of the interactome of the individual baits found a comparatively smaller number of proteins interacting with all three of them under both the states (Fig. S2A and S2B), probably due to our stringent selection criteria for the positive hits. Nevertheless, our IP sample preparation method could capture not only the DNA bound transcription complex and its interactors, but also the proteins interacting off the DNA, making the coverage specific and extensive. However, due to the largely same set of TC-interacting proteins in both the states, we focused our further analyses mainly on the active state interactome data.

**Figure 1.**
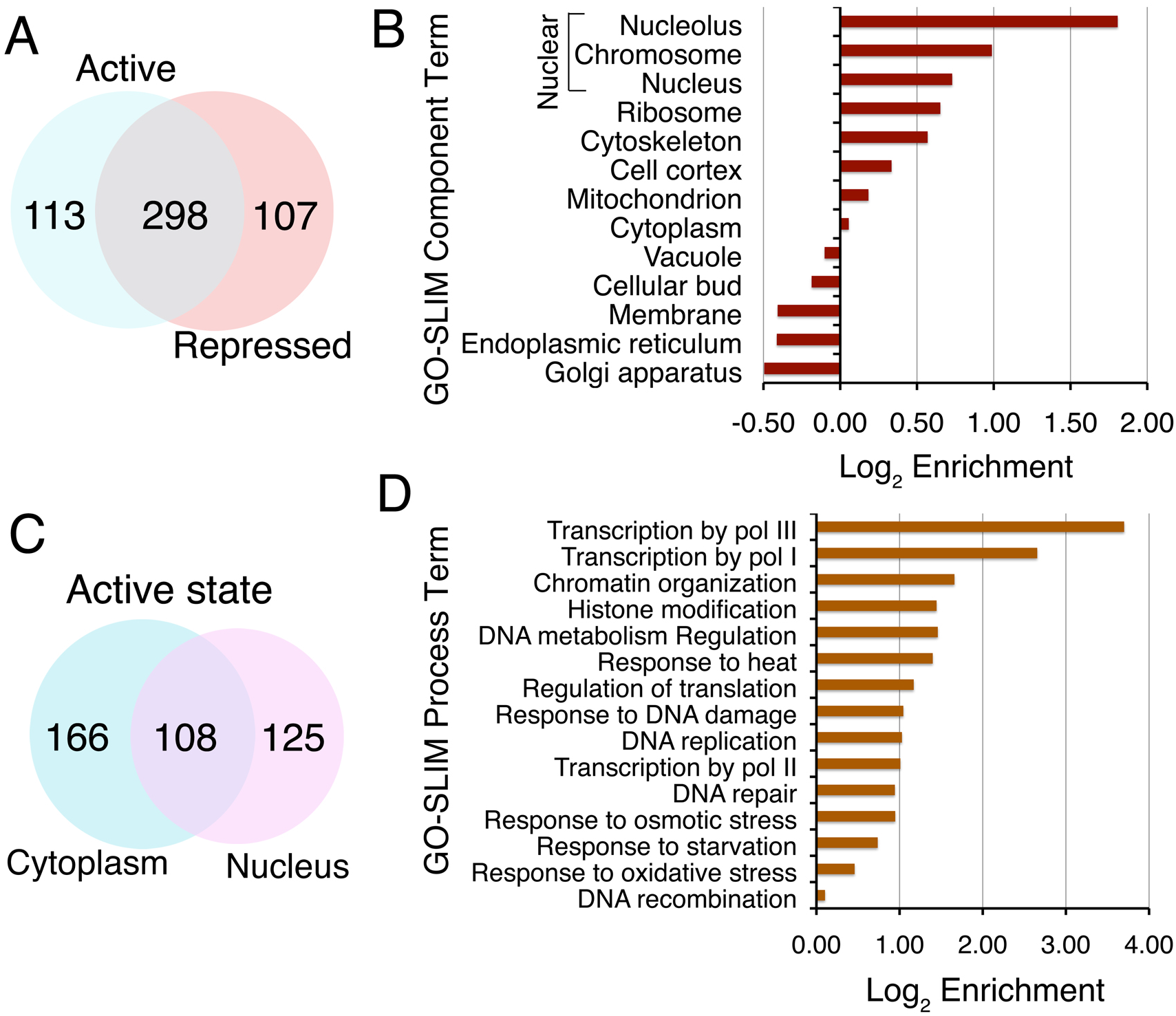
Comprehensive analysis of RNA pol III TC interactome. All the proteins confirmed as pol III TC interactors were used for various categorizations using bioinformatics analyses. Enrichment represents ratio of IP frequency to Genome frequency. IP and Genome frequencies represent the number of proteins in each category represented as % of the total number of proteins in the IP and genome respectively. (A) Venn diagram showing the overlap of pol III TC interactomes in the active and repressed states. (B) GO Slim Component mapping shows pol III TC interactome is constituted majorly by proteins from nuclear and cytoplasmic cellular fractions. (C) Venn intersections show contribution of cytoplasmic and nuclear proteins to the pol III TC interactome in the active state. (D) GO Slim Biological process analysis shows that proteins required for a variety of biological processes are enriched in the pol III TC interactome. A large fraction is comprised of proteins required for chromatin organization, and a number of chromatin modifiers.

### RNA pol III TC interactome includes cytoplasmic proteins

As the normal, functional location of a transcription complex is nuclear, GO Slim mapper component analysis revealed that a large fraction of the identified interacting proteins was nuclear, involved in a variety of nuclear processes (Fig. 1B). Nucleolar and ribosome-related proteins constitute ∼27% of the interactome (Fig. 1B), with majority of the remaining nuclear proteins being chromosomal. Interestingly, besides nuclear proteins, a significant fraction of the interactome was constituted by cytoplasmic proteins (including cytoskeletal and mitochondrial proteins, Fig. 1B). This suggests either a cytoplasmic location for pol III TC components or some interacting proteins may be shuttling between cytoplasm and the nucleus, enabling their association with TC inside the nucleus. Indeed 26% of proteins in both active and repressed states (Fig. 1C and S2C) are known to exhibit such a dual localization. They include some of the pol III (Rpc6, Rpc9 and Rpc10) and TFIIIC (Tfc3 and Tfc4) subunits classified as both nuclear and cytoplasmic in the Saccharomyces Genome Database (SGD; http://www.yeastgenome.org/). Cytoplasmic assembly of the multi-subunit pol III (51, 52) and an earlier report on human pol III activity even in the cytoplasm (53) support the possibility of cytoplasmic interactome for pol III TC.

### An extensive network of pol III TC with variety of functional complexes

Functional characterization using the GO Slim-mapper process categorization (Fig. 1D) found the pol III TC interactome rich in protein-protein interactions specific to several different biological processes in the active state. Since transcription by pol III is tightly regulated in response to growth stage, growth conditions and environmental stress (54), proteins related to different stress conditions like starvation, oxidative and osmotic stress were comparatively less enriched in the active state (Fig. 1D). Apart from the DNA metabolism and transactions, proteins related to other important processes required for accomplishing the basic physiological processes of a cell were also enriched. Proteins related to ribosome biogenesis were found at levels higher than expected according to their genome frequency (Fig S2D).

Several of the proteins and complexes unveiled in our data have been earlier reported to have genetic/physical or functional association with pol III-transcribed genes (11), which underscores the validity of our data. We used ClueGO in the Cytoscape environment to reconstruct the interaction networks, if any, of the macromolecular complexes found associated with pol III TC. This analysis revealed 27 significant complexes of which eight constitute at least 3 nodes (Fig. 2). Such interconnections of different complexes via some common partners give an intricate network of protein-protein interactions in pol III TC interactome.

**Figure 2.**
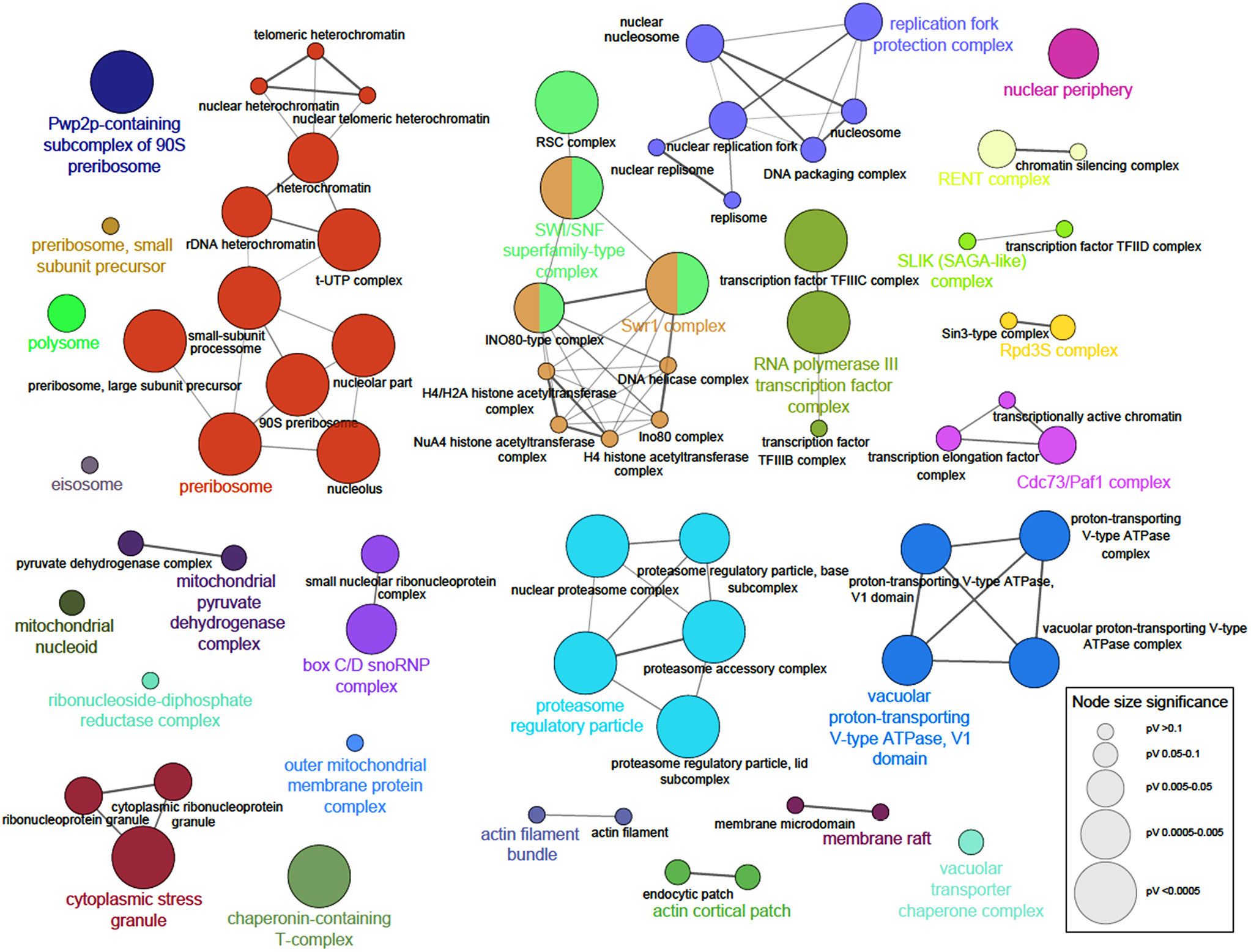
Functional complexes in the pol III TC interactome. Pol III TC interactome data were analysed to find the functional complexes enriched with pol III TC. A graphical representation of the functional complexes interacting with Pol III TC in the Cytoscape display based on the calculated p-values is shown.

### Paf1 complex associates with the pol III TC

Significantly, all five subunits of the Paf1 complex, viz. Paf1, Leo1, Ctr9, Cdc73 and Rtf1 (34, 55) were found enriched in the active and repressed states of the two (Tfc6 and Brf1) out of the three baits of the pol III TC. The Rtf1 subunit was earlier reported to nonspecifically bind FLAG M2 agarose beads (24) due to a sequence degeneracy of the amino acids 196-202 in the protein to the FLAG epitope (56). In our MS data, Rtf1 was found in the control IP of one of the three baits, Brf1 (Fig. 3A). A relatively very low number of SSCs for the functionally similar subunit Leo1 (34) were also found only in the Brf1 control IP. No other Paf1 subunit was found in any other control. This data indicated that while Paf1 complex interacts with TFIIIC and TFIIIB in both the states, interaction with pol III is favored in the repressed state. We found that in reciprocal pull-downs the Paf1 subunit co-immuno-precipitates equally well with all three components of the pol III TC in active or repressed states (Fig. 3B-E and Fig.S3), suggesting association of Paf1 with TC may not be related to the active transcription process.

**Figure 3.**
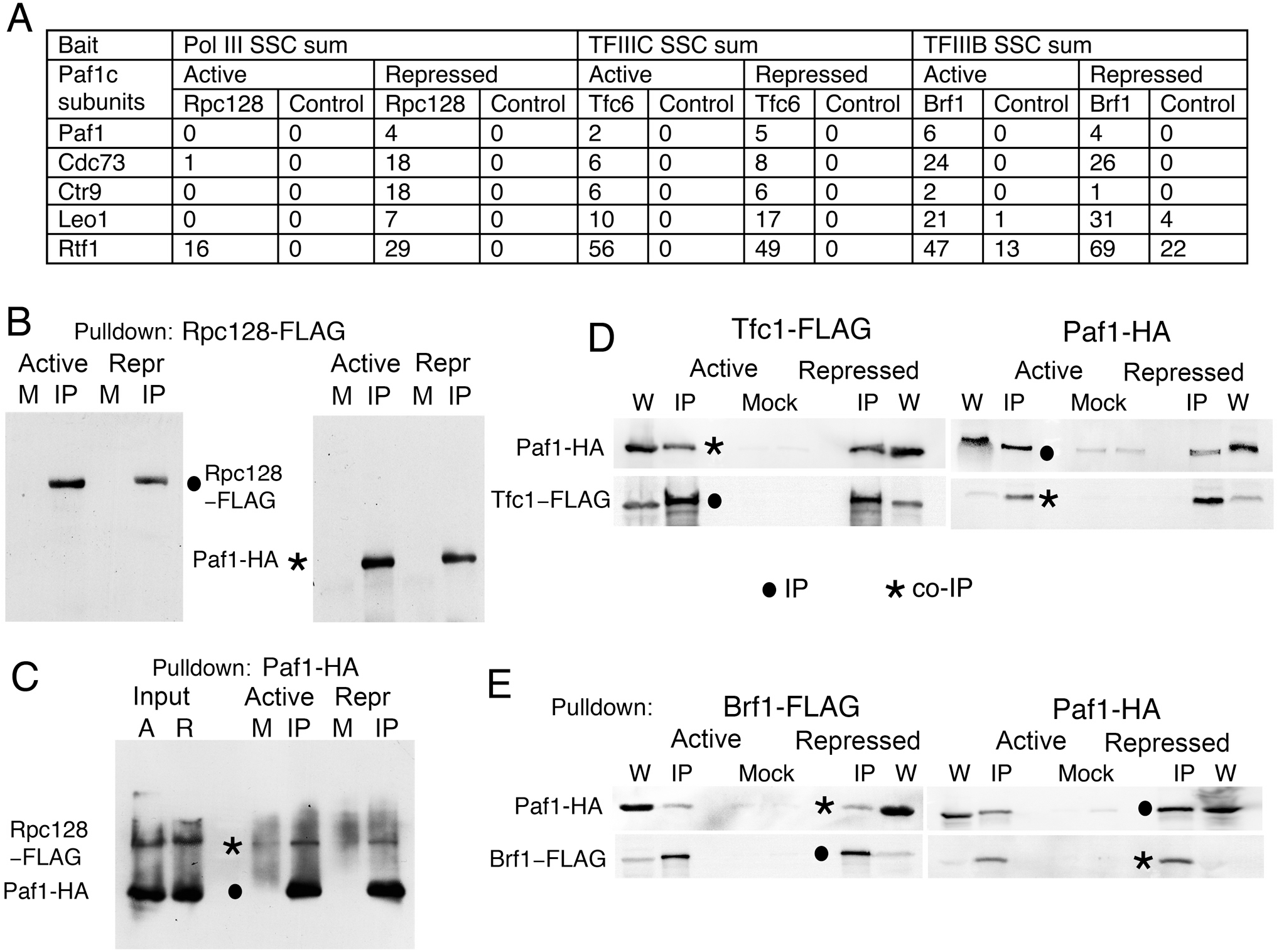
Paf1 associates with pol III TC in active and repressed conditions. (A) Data from AP-MS/MS on Paf1 complex and pol III TC interaction. Sum of SSCs (Specific Spectral Counts) for each subunit of Paf1 complex in the IP and control samples of the three baits is given. (B)-(E) The reciprocal co-immunoprecipitations show interaction of the Paf1 subunit with the pol III TC. Lanes M: Mock, IP: Immunopulldown, W: whole cell extract. IP and co-IP bands are marked in every panel with a dot and asterisk respectively. Paf1 interacts with Rpc128 (B) and (C), Tfc1 (D) and Brf1 (E) in both active and repressed conditions. (B) and (C) Full blots are shown for the co-IPs performed by using Rpc128-Flag (B) or Paf1-HA (C) as bait and blots were sequentially probed with the antibodies one at a time. Repr and [R] denote repressed while [A] stands for active state. (B) The blot was stripped and washed between the first and second sequential probings. (C) Blot of Rpc128 and Paf1 co-IP was not stripped before second probing with the anti-Flag antibody. The first probing showing Paf1-HA IP is shown in the Fig. S3A. (D) and (E) Sequential probings of the blots with anti-Flag and anti-HA antibodies, one at a time. Blots were completely stripped before the second probing. Input signal with Flag probing was generally found low in these probings. The bands are shown as cropped images for the purpose of clarity. Full blots without intermediate steps of strippings are shown in the Figures S3B-D.

### Paf1 complex associates with the pol III-transcribed genes

The presence of Paf1 complex in the pol III TC interactome was intriguing. In order to ascertain whether this interaction is complemented by the PAF1C presence on the pol III-transcribed genes in vivo; we analysed two sets of the earlier published and available genome-wide occupancy data (45, 46). The genome-wide Rtf1 occupancy data (46) showed its diverse but low levels at individual genes (Fig. S4A). As compared to pol II-transcribed genes, the average occupancy of Rtf1 was found lower on the pol III-transcribed genes (Fig. S4B). Reanalysis of the data after filtering out the tRNA genes with pol II-transcribed gene ends within 300 bp on their both the sides did not change the average profile of Rtf1 occupancy on the tRNA gene region (Fig. S4B). Therefore, the observed occupancy on the tRNA genes is not due to the vicinity of pol II-transcribed genes (Fig S4B). Further analysis of an earlier ChIP-exo, genome-wide occupancy data (45) found highly similar occupancy profiles of all five subunits of the Paf1 complex (Fig. 4A and S4C). The average occupancy profiles of all five Paf1C subunits show lower levels on the gene body and a sharp peak at the −10 bp position between the box A at +20 bp position (22) and the biding site of the pol III transcription initiation factor TFIIIB at the −30 bp position (Fig. 4A and S4C). As subunits of the same complex, Ctr9 occupancy on the individual genes (Fig. S4D) showed a positive correlation coefficient with both Paf1 and Rtf1 occupancies (r = 0.414 and 0.471 respectively).

**Figure 4.**
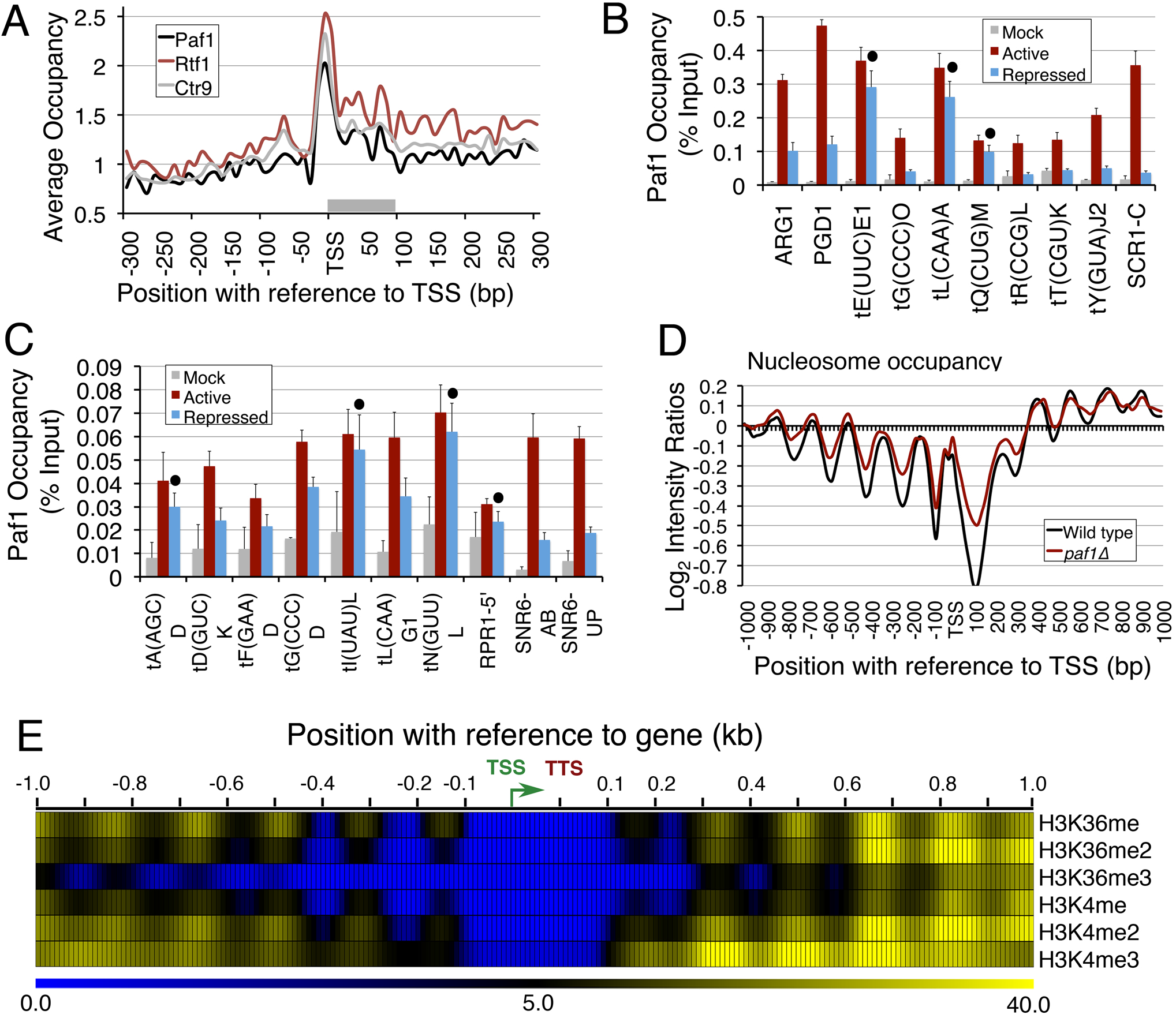
Paf1 localizes to pol III-transcribed genes. Low level of Paf1 occupancy is found at the pol III-transcribed gene loci. (A) and (B) The genome-wide data on Paf1 occupancy (45) was analysed. Similar average occupancy profiles were found for the subunits on the tRNA genes. The average occupancies on both 300 bp upstream and downstream of TSS are plotted. The grey bar marks the tRNA gene region on the X-axis. (B) and (C) Paf1 occupancy over pol III-transcribed genes relative to TelVIR region were estimated by ChIP and Real time PCR in active and repressed states. The changes were found to be significant (p value <0.04) for all except those marked with a dot. (D) Paf1 does not influence nucleosome occupancy (48) on tRNA genes. (E) Methylation levels on K4 and K36 residues of the histone H3 (49) are very low near tRNA genes. Heat map is shown for 1 kb upstream and downstream of the gene. Color gradient code is shown at the bottom, TTS and TSS (bent arrow) are marked.

Using pol II-transcribed *ARG1* and *PGD1* as positive controls, we checked for the presence of Paf1 on several pol III-transcribed genes by the ChIP-Real Time PCR assay (Fig. 4B and 4C). We found Paf1 on all of the tested tRNA genes, which could be grouped into comparatively higher (Fig. 4B) or lower (Fig. 4C) occupancy level genes. As compared to the *ARG1* gene, Paf1 level was ∼2-3 fold lower on the high (Fig. 4B) and >3 fold lower on the low occupancy group genes (Fig. 4C). The non-tRNA genes *SNR6, RPR1* show low Paf1 occupancy (Fig. 4C) while *SCR1* (Fig. 4B) could be grouped with higher occupancy genes (amplicons shown in the Fig. S4E). Interestingly, under repression, ten of the total seventeen tested pol III-transcribed genes show loss while rest of the genes do not show any significant change. Thus, similar to pol II-transcribed genes (29, 57), Paf1 may be involved in both positive and negative regulation of the pol III-transcribed genes in a gene-specific manner.

Since Rtf1 is known to be a part of the PAF1 complex in yeast (55, 56), the presence of rest of the subunits in our IP-MS data as positives, the Paf1 interaction with the pol III TC in both active and repressed states (Fig. 3), presence of all five subunits on the tRNA genes in active state and loss of Paf1 in the repressed state (Fig, 4) affirm the interaction of pol III TC with the complete Paf1 complex, which may be linked to the transcription status of the genes and not to the steps of the transcription process.

### Paf1 does not affect nucleosomes near the pol III-transcribed genes

Paf1 is reported to influence nucleosome occupancy on pol II-transcribed gene regions (58–60). As transcription on pol III-transcribed genes is modulated by the gene-flanking positioned nucleosomes (61) and Paf1 is found on the pol III-transcribed loci (Fig. 3), we retrieved and analysed the published data on the nucleosome occupancy on these genes in wild type and *paf1*Δ cells (48). Occupancy of the gene-flanking nucleosomes shows only a small change in the *paf1*Δ cells (Fig. 4D). Therefore, Paf1 does not influence positions or occupancies of gene-flanking nucleosomes on the tRNA genes.

### Low histone methylations on the pol III-transcribed genes

Paf1 is known to mediate repression of a pol II-transcribed gene *ARG1* by preventing Gcn4 recruitment to the *ARG1* promoter and subsequent histone H3 acetylation (62). The Paf1 complex is required for recruitment of Set1 and Set2 methyltransferases on pol II-transcribed gene bodies during elongation (32, 63). Set1 and Set2 mark active genes with H3K4 and H3K36 methylations, respectively (64, 65). Unlike the earlier reported presence of active chromatin marks near human pol III-transcribed genes (66–68), our analysis of the published genome-wide data on histone modifications (49) in yeast revealed absence/extremely low levels of H3K4 and H3K36 methylations, upstream of TSS on the tRNA genes (Fig. 4E).

As compared to the genome-wide levels, nucleosome occupancy around tRNA genes is known to be low (69, 70). Therefore, low levels of H3K4 and H3K36 methylations found on pol III-transcribed tRNA genes may be due to low nucleosome occupancy and/or low Paf1 levels on these gene regions. Further validations and measurements of these methylations in *paf1*Δ cells will be required to get a better understanding of Paf1 role in histone methylations on different pol III-transcribed genes. Nevertheless, in the light of the short transcribed regions and nucleosome-free gene bodies, the significance of the association of Paf1 complex with pol III-transcribed genes may be different from its known roles on pol II-transcribed genes.

### Genotoxin MMS effect on the pol III-transcribed genes is Paf1-independent

Cells lacking Paf1 complex subunits show different genotoxin sensitivities (71, 72). The tRNA genes are prone to the DNA damage in vivo due to the slowing down of replication on them (6, 7, 73). Genome-wide mapping of γ-H2A enrichment has been reported to represent the fragile as well as replication fork stalling sites in the genome (73). We found low levels of γ-H2A on the pol III-transcribed genes in the wild type cells under normal growth conditions (Fig. S5A), which represent the slow replication on these highly transcribed genes. The γ-H2A levels at individual genes in the *paf1*Δ cells were found at levels higher than the wild type cells (Fig. S5B), which suggests gene-specific increase of DNA damage in the *paf1*Δ cells, As compared to the wild type cells, higher γ-H2A levels were found in the *paf1*Δ cells, when both wild type and mutant cells were similarly exposed to the genotoxins HU (Fig. 5A) or MMS (Fig. 5B). The increase in γ-H2A levels in the *paf1*Δ cells was more pronounced under HU (2-8.5 fold, p<0.005) than under MMS (1.3-1.8 fold, p<0.05) exposure, varying in a gene-specific manner (Fig. S5C). This is consistent with the earlier observed higher sensitivity of the *paf1*Δ cells to HU than to MMS (71, 72).

**Figure 5.**
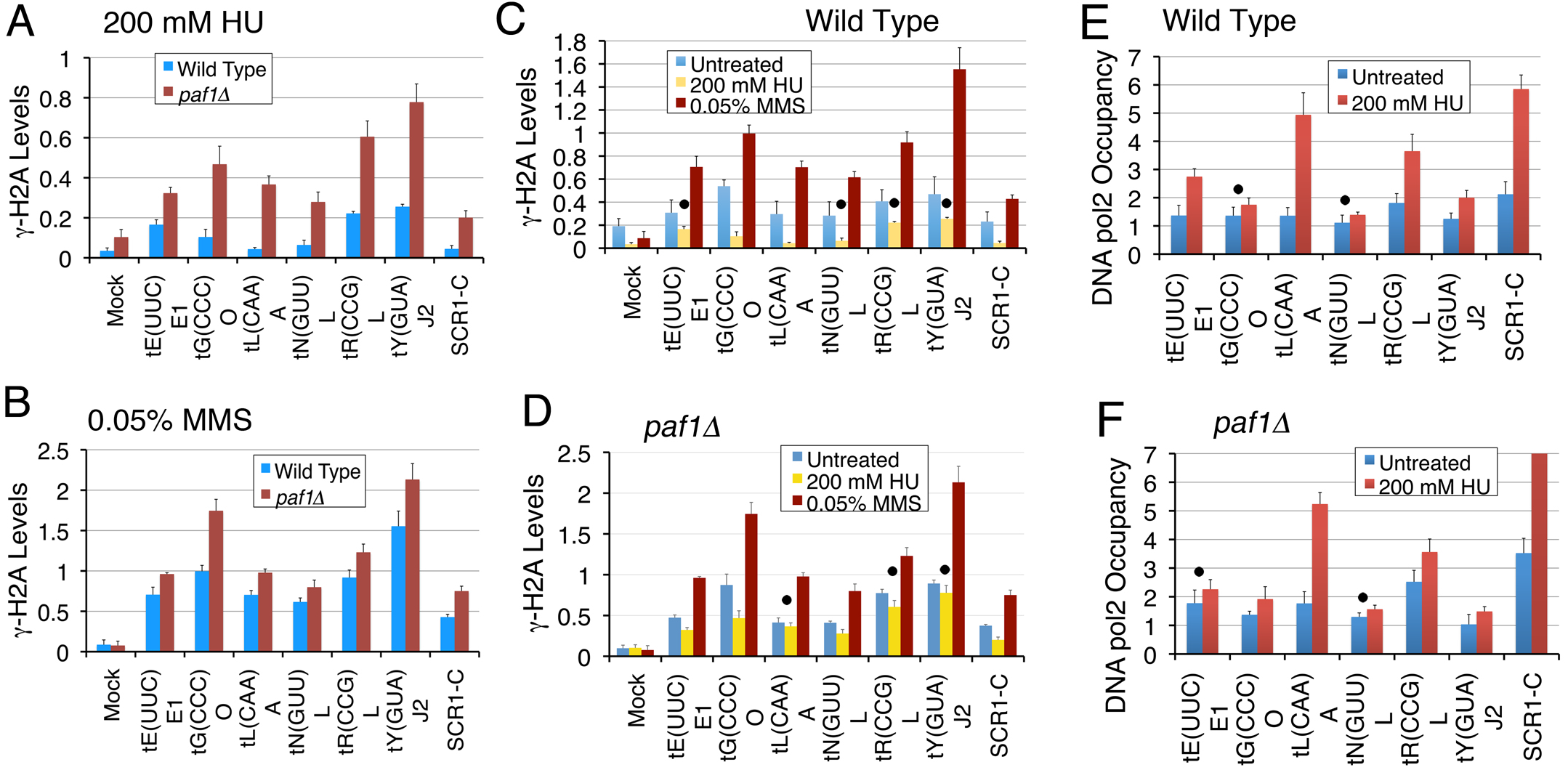
Paf1 is required to counter the replication stress at pol III-transcribed genes. Levels were measured in cells with or without exposure to genotoxins by using ChIP and Real Time PCR method. Dots mark the genes, which do not show significant change under a specific condition (p value <0.05). Rest of the changes are significant, (A) and (B) As compared to wild type, γ-H2A levels increase in *paf1Δ* cells when both type of cells are similarly treated with (A) HU; all p values <0.0045 or (B) MMS; all p values <0.05. (C) and (D) As compared to the levels in the untreated condition, the γ-H2A levels increase (p value <0.009) after MMS exposure. (C) As compared to the low, normal (untreated) levels in the wild type cells, γ-H2A levels decrease when exposed to HU. The p values for the genes marked with a dot vary between 0.06 and 0.076; rest are <0.035. (D) Higher than the wild type γ-H2A levels in the *paf1Δ* cells mostly do not change when exposed to HU. The p values for the genes marked with a dot vary between 0.05 and 0.16; rest are <0.04. (E) and (F) The DNA pol2-9XMyc occupancy on the tRNA genes increases upon exposure to 200 mM HU for 2hr. (E) In the wild type cells, DNA pol2 shows gene-specific increases on most of the genes, except *tG(CCC)O* (p value = 0.061 and *tN(GUU)L* (p value = 0.052), marked with a dot. (F) HU treatment of the *paf1Δ* cells brings the DNA pol 2 levels on all the tested genes to same as that in the wild type cells, except on the *SCR1* gene where levels were found beyond the given scale (increase of ∼6.8 fold).

Genotoxins HU and MMS are known to cause replication stress/DNA damage via different mechanisms. As opposed to untreated controls, exposure to MMS (causes DNA damage via bases alkylation) resulted in an increase of the γ-H2A levels in both wild type and *paf1*Δ cells as expected (Fig. 5C and 5D). Surprisingly, as compared to untreated controls, the γ-H2A levels upon HU exposure (inhibits replication via dNTPs depletion) show either a loss (mostly wild type cells) or no change (mostly *paf1*Δ cells) in a gene-specific manner (Fig. 5C and 5D). These data show that Paf1 has a greater role in countering HU toxicity as compared to MMS toxicity on the pol III-transcribed tRNA genes.

### Paf1 counters the replication stress at the tRNA genes in a gene-specific manner

Loss of γ-H2A is reported to signify DNA repair at low levels of damage (74). In the untreated wild type cells, low γ-H2A levels (Fig S5A) signify only a minimal and gene-specific DNA damage at the pol III-transcribed genes. Exposure of the wild type cells to HU inhibits replication and represses pol III transcription (75). Hence reduced replication and transcription upon HU exposure may enable repair and reversal of the pre-existing low-level DNA damage (Fig. S5A), which results in observed loss of the γ-H2A levels (Fig. 5C). With the slow replication and replication fork pausing, the DNA polymerase may be found at the genes in the normally cycling cells. An earlier, low-resolution microarray study had found enrichment of DNA pol2 at highly transcribed genomic sites, including three tRNA genes in the unsynchronized wild type cells (76). Consistent with this, our ChIP measurements found only low DNA pol2 levels in the normally cycling cells, which increased on most of the tested genes after exposure to 200 mM HU, in a gene-specific manner (Fig. 5E).

In the *paf1*Δ cells, the γ-H2A levels on most of the genes did not show loss upon HU-exposure (Fig 5D), probably because the reported 2 fold decrease of *RNR1* expression may cause additional replication stress on the genes in the untreated *paf1*Δ cells (57, 71). Consistent with this, in the absence of any genotoxin, the pol2 levels on different genes in the *paf1*Δ cells were found at levels either similar to or higher than the wild type levels (Fig. S5D). Moreover, the pol2 increase on each gene in *paf1*Δ cells was only half of the increase with HU exposure in the wild type cells (cf. Figs. 5E and S5D). In comparison, excepting *SCR1*, the pol2 levels in the *paf1*Δ cells increase to almost the same level as seen in the wild type cells upon HU exposure (cf. Figs. 5E and 5F). These results suggest that influences of Paf1 and HU on the replication stress on the tRNA genes are opposite and Paf1 plays a damage-preventive role on the tRNA genes during the replication fork progression. The observed increase of γ-H2A levels in *paf1*Δ cells, due to the chemically induced DNA damage by MMS (Fig 5B) matches with this conclusion.

### Pol III occupancy decreases upon replication stress

Exposure of the wild type cells to HU results in quick eviction of pol II from the replication-transcription encounter sites at pol II-transcribed genes (72). Unlike pol II-transcribed genes (72), the Paf1 occupancy on the tested pol III-transcribed genes does not change (Fig. 6A) when wild type cells are exposed to the genotoxins. High transcription at the pol III-transcribed genes is proposed to interfere with smooth progression of replication machinery (6). Therefore, we estimated pol III levels on the genes when exposed to genotoxins. Consistent with earlier reported repression of the tRNA gene transcription under replication stress (75), the pol III levels dropped significantly when wild type cells were exposed to genotoxins HU or MMS (Figs. 6B and S6A respectively). Reducing the pol III occupancy may be a cellular response to the replication stress caused by the genotoxins in order to reduce the transcription rate and facilitate the replication fork progression on the tRNA genes. Similar to wild type cells, the pol III levels drop in the genotoxin-sensitive *paf1*Δ cells when exposed to 0.05% MMS (Fig S6B).

**Figure 6.**
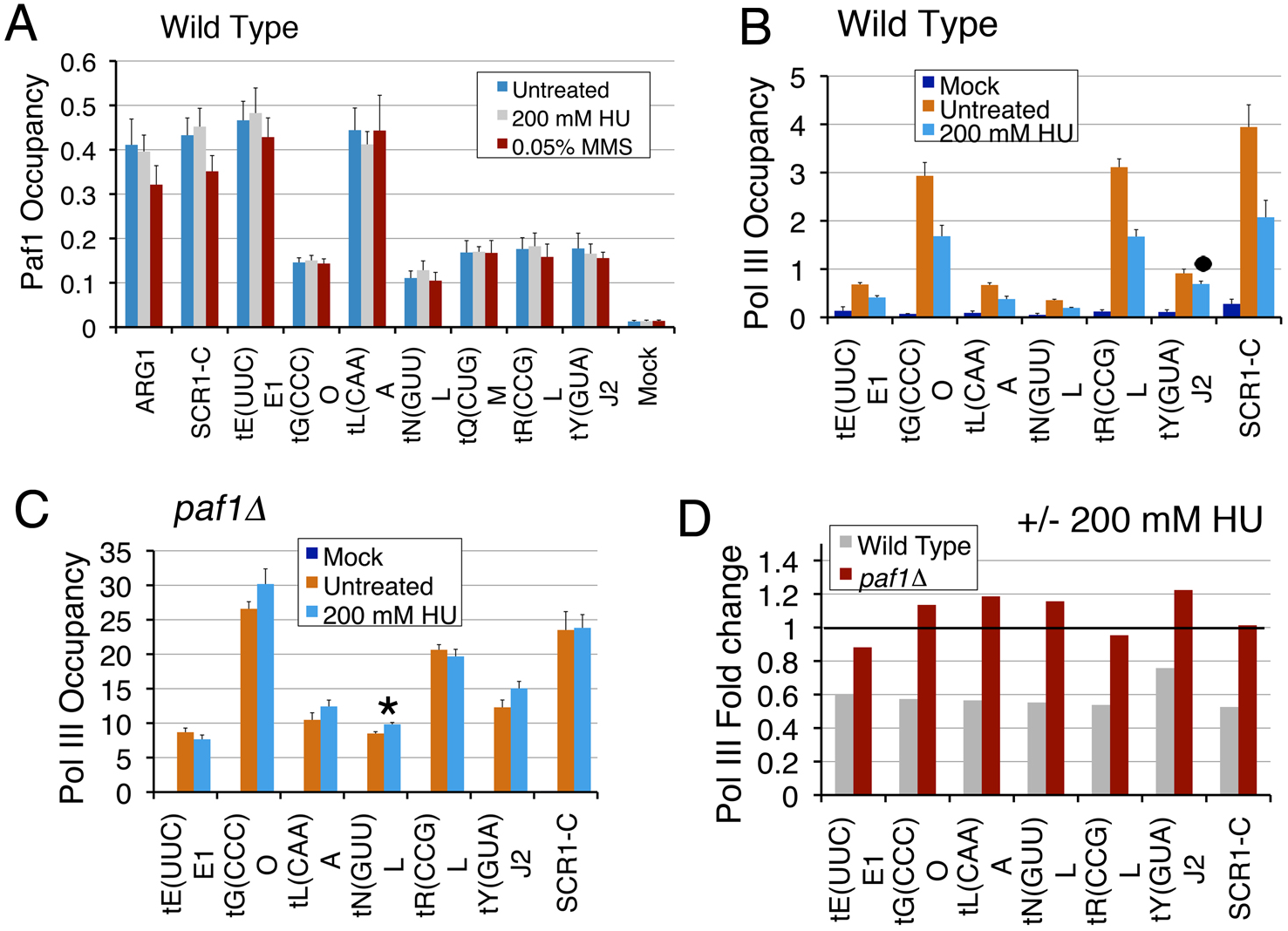
Pol III is lost from the genes under replication stress in a gene-specific manner. (A) Paf1 occupancy on the tested genes in the wild type cells does not change when exposed to the genotoxins HU or MMS. (B) Pol III occupancies in normally cycling wild type cells decrease upon exposure to HU. The dot marks the gene with insignificant decrease (p value = 0.29). (C) In the *paf1Δ* cells, pol III occupancies are not affected upon exposure to HU, except on *tN(GUU)L* (p value= 0.045), marked with an asterisk. (D) Data in the panels B for wild type and panel C for the *paf1Δ* cells were used to obtain the ratios HU treated/untreated cells. As a value of 1.0 (marked with a horizontal line) denotes no change, ratios on the y axis reveal that HU causes ∼40% loss of pol III in the wild type but no change in the *paf1Δ* cells.

In contrast to MMS effect, the pol III levels in the *paf1*Δ cells do not decrease or alter on exposure to HU (Fig. 6C). We compared the fold decrease in pol III with genotoxin exposure by calculating the ratios of the corresponding pol III levels on each gene in the *paf1*Δ and wild type cells (Fig. 6D and S6C). Consistent with the results so far, similar pol III decrease for both types of cells is found when exposed to MMS (Fig, S6C), confirming MMS toxicity is not Paf1-dependent. In comparison, HU exposure affects pol III levels more severely in the *paf1*Δ as compared to the wild type cells (Fig 6D). We conclude that Paf1 is required to counter the HU-induced replication stress on the pol III-transcribed genes.

### Pol III occupancy is highly increased upon Paf1 deletion

Human Paf1 is reported to regulate the release of paused pol II levels at the 5’ gene-ends (36, 77, 78). We found the yeast pol III level was highly increased on all the tested genes in the *paf1*Δ and all but three of the tested genes in the *rtf1*Δ yeast cells (Fig. 7A, 7B and S7A). The total pol III levels (Rpc128 subunit) remain at the wild type levels in the *paf1*Δ and *rtf1*Δ cells (Fig. 7C). The presence of pol III TC components on the tRNA gene bodies protects them against MNase digestions (23, 79). Therefore, the increase in pol III levels at the gene body may be the reason for the apparent increase in intensity on the gene body in the MNase digestion profile of the chromatin form the *paf1*Δ cells (Fig, 4D). The high increase also explains why no loss in pol III levels are seen upon HU exposure in the *paf1*Δ cells (Fig. 6C).

**Figure 7.**
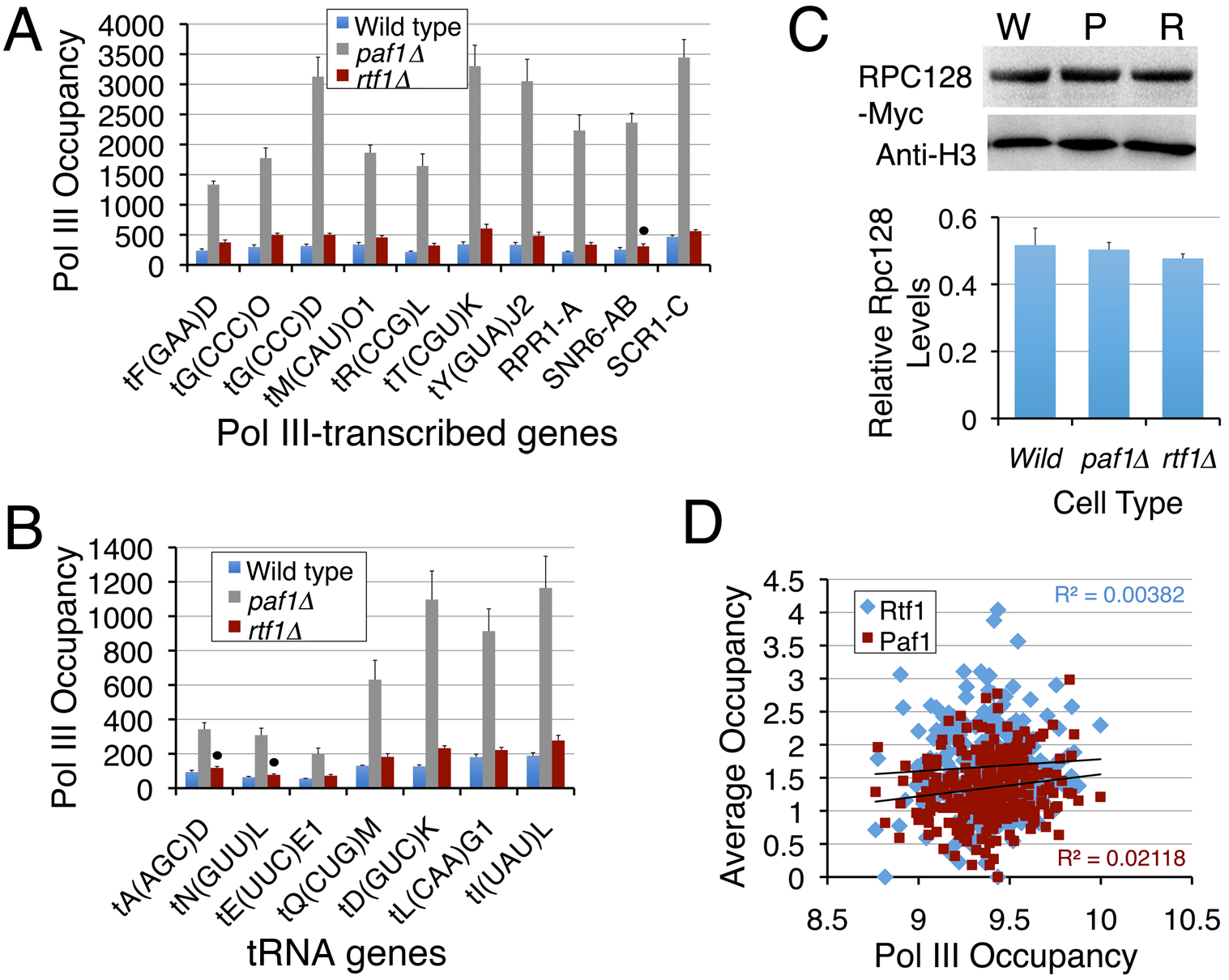
Pol III occupancy shows gene-specific increases in the *paf1Δ* and *rtf1Δ* cells. (A) and (B) Rpc128 occupancy at pol III-transcribed genes relative to TelVIR region in wild type, *rtf1Δ* and *paf1Δ* cells. Averages and scatter from three biological replicates are plotted. The dot denotes change is non-significant in *rtf1*Δ cells, probably due to higher scatter. (C) Western quantification of the total levels of Rpc128 protein in the wild type (W), *PAF1* (P) or *RTF1* (R) deletion cells. Lower panel shows the Rpc128 protein levels normalized against H3 do not differ in the three types of cells. (D) Paf1, Rtf1 (45) and pol III (61) occupancies on the tRNA genes are compared. Occupancies of Paf1 and Rtf1 (at −205 to 155 bp) do not show any correlation with pol III levels (at −200 to 300 bp).

The pol III occupancy is affected (Figs. 7A and 7B) more by Paf1 deletion (∼3.5-10.2 fold increase) than by the Rtf1 deletion (∼1.2-1.8 fold increase). The Paf1 (and not Rtf1) deletion affects protein levels of other Paf1C subunits as well (80). This suggests that the *PAF1* deletion effects may be more severe than the *RTF1* deletion effects. In agreement, as compared to Paf1, Rtf1 was earlier found to show only a minor repressive effect on pol II-transcribed *SER3* expression (59). Rtf1 shows unequal distribution (Fig. S4A) while similar pol III occupancy is found on the individual tRNA genes (61) in the wild type cells. A comparison of average Paf1 and Rtf1 levels on the genes showed only a poor correlation with the pol III levels (Fig. 7D), although the correlation is better for Paf1 (R= 0.1455) than for Rtf1 (R= 0.0618). Consistent with these analyses, the pol III occupancy on different genes is elevated to different levels in the *paf1*Δ cells (Figs. 7A and 7B), indicating a gene-specific variation in Paf1 influence on the pol III levels.

### Paf1 has a gene-specific influence on the transcription of tRNA genes

Paf1 has been shown to promote transcription of pol II-transcribed genes through chromatin in vitro (35). Genome-wide studies have reported an unequal effect of Paf1 deletion on expression of the Paf1-associated pol II-transcribed genes (29, 57), suggesting a role for it in transcription of only a limited number of genes. Paf1 is reported to have a repressive role in the expression of pol II-transcribed *SER3* and *ARG1* genes (59, 62). Our measurements of transcription in wild type and Paf1 or Rtf1 deletion cells by northern probing did not find any change in levels of 18S rRNA (pol I transcript), U4 (pol II transcript) and U6 snRNAs and 5S rRNA (Fig. 8A). Since the pre-tRNAs (primary transcripts) are generally difficult to detect due to their low levels as compared to mature tRNA levels, we could measure only three pre-tRNA levels by these estimations (Fig. 8B). Nevertheless, as compared to wild type, both primary (p<0.05) and mature (p<0.03) tRNA levels showed a small but significant increase for the tested genes in both the deletion strains (Fig. 8C and 8D). Although measurement of primary transcript from a larger number of genes may be required, above results suggest a repressive role for PAF1C in the pol III transcription, which could be related to a role of Paf1 in increasing pol III pause in the wild type cells (Fig. 7).

**Figure 8.**
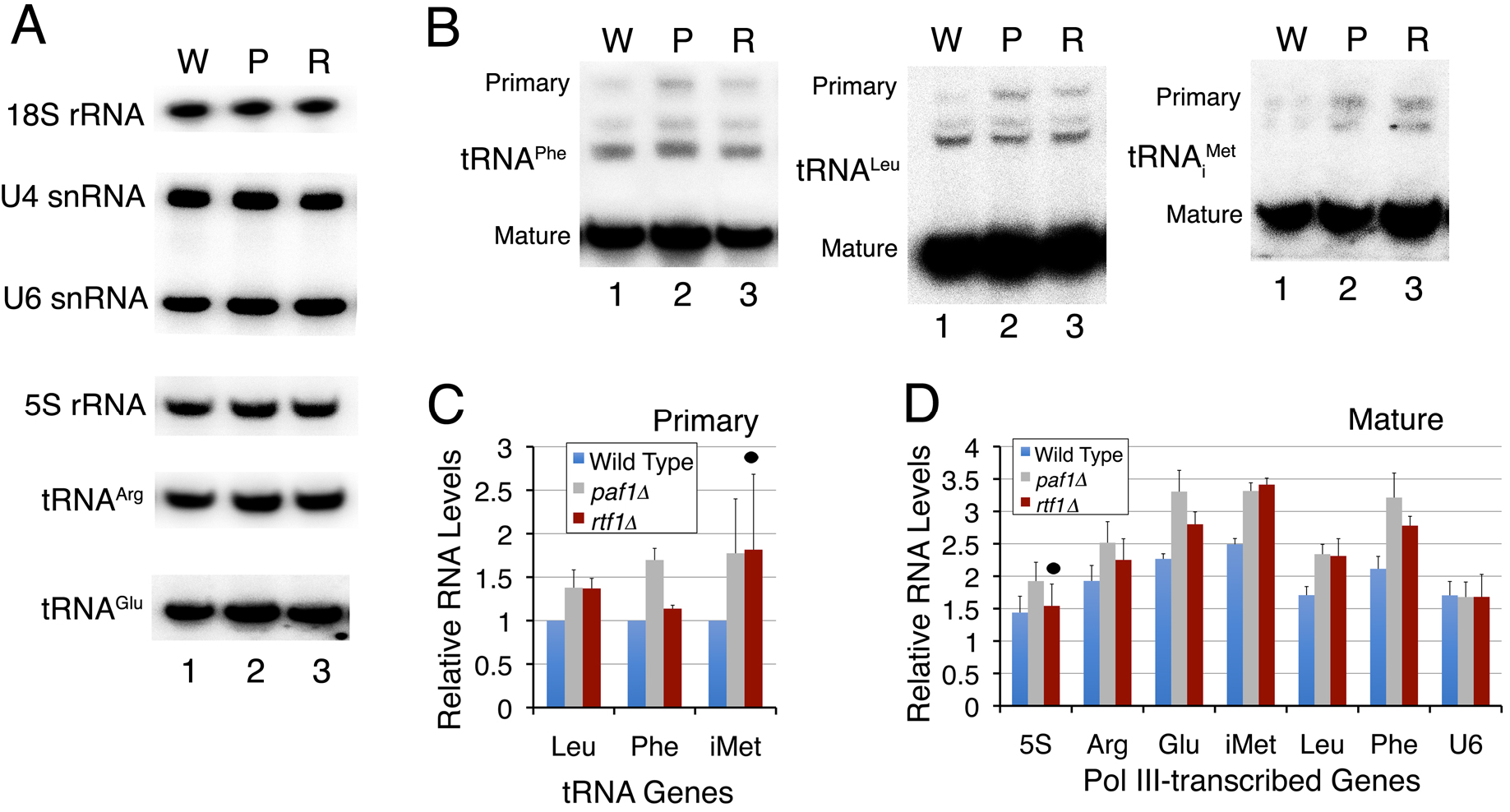
Paf1 has repressive role in RNA pol III transcription. (A) and (B) Estimation of transcript levels in *paf1Δ* and *rtf1Δ* cells by Northern analysis for different genes in wild type [W] *paf1Δ* [P] and *rtf1Δ* [R] cells. Radiolabeled oligonucleotide probes individually against 18S (pol I transcribed), U4 (pol II transcribed) and pol III transcribed U6, 5S, different tRNA genes were used individually. Primary and mature transcript level positions are marked. (C) Quantification of pre-tRNA products in panel A for tRNA_i_^Met^, tRNA^Phe^ and tRNA^Leu^ normalized with U4 levels. Averages and scatter from three independent experiments are plotted. (D) Quantification of mature RNA products in panel A for 5S rRNA and tRNA_i_^Met^, tRNA^Arg^, tRNA^Phe^ and tRNA^Leu^ normalized with U4 levels. Average and scatter from three independent experiments is plotted. Dots mark the statistically insignificant changes.

Most of the tRNAs are represented by multiple isogenes, which make their estimations by using gene body probes, erroneous. Therefore, pol III occupancy is considered a better measure of the transcription activity of these genes. Our estimations show that the high increase of pol III levels in *paf1*Δ cells does not translate into a similar increase in tRNA levels. Similar to this, an earlier observed increase/no change in pol I levels showed a decrease in the transcript levels (33). Only small (although significant) changes in transcript levels due to *PAF1* or *RTF1* deletions in this study are found consistent with lack of a correlation between Rtf1 or Paf1 occupancies with the recently published levels of pol III (Fig. S7B) bound to the presumably primary transcripts of all the tRNA genes (81). The same is expected from the poor correlation (R=0.1547; Fig. S7C) found between the pol III levels bound to the genomic sites (61) and to the primary transcripts (81; Fig. 7D). Thus, it is not surprising that the high increase in pol III levels (Figs. 7A and 7B) is not matched by a similar increase in transcription (Fig. 8) in the *paf1*Δ and *rtf1*Δ cells.

The above results suggest that the observed effects of Paf1 or Rtf1 deletions on the pol III-transcribed genes are gene-specific. Despite a highly increased pol III occupancy (Fig. 7A), no change in RNA levels in *paf1*Δ and *rtf1*Δ cells (Fig. 8A), suggests no role for Paf1 in *SNR6* transcription. Similarly, one tRNA gene *tE(UUC)E1* shows highest Paf1 (Fig. 4B) and lowest pol III (Fig. 7B) occupancy among all the tested tRNA genes. But changes in the pol III levels in the *paf1*Δ and *rtf1*Δ cells (Fig. 7B) and Paf1 level under repression (Fig. 4B) are least on the *tE(UUC)El* gene. It may be interesting to note in this connection that loss of Paf1 under starvation (Fig. 4) is not associated with increase of pol III levels as pol III is lost from all the genes under starvation (19, 61).

In summary, this study has found association of many protein complexes with the pol III TC, which are so far not reported to influence the transcription by pol III. We found one of the known pol II-associated factors, the PAF1 complex occupies the tRNA genes in gene-specific manner and associates with the pol III TC in both active and repressed states. Most of its known roles on pol II-transcribed genes are not found on the pol III-transcribed genes. The occupancies of Paf1/Rtf1 and pol III are not correlated. Yet, the pol III occupancy on the genes in the Paf1/Rtf1 deletion mutants shows high increase. This accumulation of pol III on the gene body causes steric obstruction of the transcription and replication in the *paf1*Δ cells. Accordingly, as compared to the wild type levels, only a small gene-specific increase in the tRNA levels is found in the Paf1/Rtf1 deletion cells. Pol III levels on the genes reduce upon genotoxins exposure while the low levels of DNA pol2 and γ-H2A on the individual genes in the unsynchronized wild type cells increase upon genotoxins exposure and in the *paf1*Δ cells. Taking together, the study shows a protective role of Paf1 on pol III-transcribed genes, which are under replication stress even in the normally cycling cells.

## Discussion

### A wide protein-protein interaction network of the pol III TC

The topology of the pol III TC shows that TFIIIB is found only at the 5’ end, TFIIIC on the whole gene body and pol III more towards the 5’ end and the transcribed gene region (Fig. S7A). These TC components are juxtaposed but spread in space over a short length of ∼200 bp (22, 23). Therefore, using the baits spread over the whole unit of a pol III-transcribed gene we could capture probably the entire pol III TC interactome. These proteins associate with the TC components specifically and differentially and change the pol III TC interactomes temporally and spatially according to the bait in an asynchronous culture. They may provide different signaling cues to the TC, resulting in modulation of pol III TC activity in different ways. A comparatively small number of overlapping interactors of three baits and detection of a large number of cytoplasmic proteins suggests that the pol III does not exist as a holoenzyme in the cytoplasm. One unusual prefoldin Bud27 was first found by proteomic approaches as a protein associating with the common subunit of the three RNA polymerases, Rpb5 (82). Bud27 was initially found necessary for the assembly of all the three polymerases (83). Later it was found to influence Rpc160/128 protein levels and pol III biogenesis in the cytoplasm (52). Not surprisingly, out of the three baits, Bud27 is found to interact with only pol III in our data.

### Rtf1 as part of the Paf1 complex associated with the pol III TC

Paf1 has been reported to associate with and influence pol I and pol II transcription (33, 34, 55, 56). This is the first study reporting Paf1 association with pol III TC and its influence on the replication stress at the pol III-transcribed genes. One of the subunits Rtf1 has been found to show less stable association with the PAF1C in all species other than the budding yeast (84–86). Requirement of yeast Rtf1 for recruitment of PAF1C on the pol II-transcribed genes (87) suggests it may be located peripherally in the complex. Pull down of only Rtf1 with pol III in the active state, possibly as the anchor subunit of the PAF1C (30, 87), suggests a weak contact of Rtf1 with rest of the PAF1C subunits. Thus, during transcription in the active state, highly dynamic state of pol III may induce dissociation of non-Rtf1 subunits from pol III. However, taking together, our other estimations show that Rtf1 is part of the PAF1C on the pol III-transcribed genes as well. Several studies in the past have found that Rtf1 does not behave similar to other subunits despite being part of the same complex in yeast (34, 56, 71, 72, 84). Similarly, the deletion of the Rtf1 or Paf1 results in increase of pol III to different levels in this study.

### PAF1 complex influences pol III-transcribed gene loci in a gene-specific manner

The pol III-transcribed genes generally do not follow a single, common pattern of response under a condition and most of the times respond in a gene-specific manner (79). A recent study has demonstrated that Paf1 levels on pol II-transcribed genes vary independent of its role in elongation (88). Paf1 levels higher than a low, basal optimum required for transcription elongation, serve regulatory role whereby the gene-specific variations decide the fate of the transcripts (88). By analogy, differences in the Paf1 levels may influence the transcription in a gene-specific manner on different pol III-transcribed genes. Similar to their reported behaviour in pol I and pol II transcription (33, 56), the deletion of Paf1 subunits shows different effects on transcript levels and pol III occupancy on different genes, tested in this study. Paf1 loss under starvation from most of the high but not low occupancy pol III-transcribed genes implies higher gene-specific Paf1 association under normal growth conditions.

### Recruitment of PAF1 complex near TSS causes pol III pausing

Increased pol III levels on the tRNA genes may represent even increased recruitment. Higher pol III recruitment on the genes may increase transcription while stalling represents reduced transcription. Our results showing only a small change in the tRNA levels with high increase of pol III are consistent with pol III stalling rather than increased recruitment in the *paf1*Δ and *rtf1*Δ cells. Further, a lack of correlation in occupancies of Paf1 and Rtf1 with pol III suggests PAF1C is not recruited by pol III. Its interaction with TFIIIC and TFIIIB in both active and repressed states and peak occupancy at the −10 bp position suggest it is recruited to the genes by TFIIIB or TFIIIC probably before or simultaneous with the joining of pol III into the TC.

The earlier reported average occupancy profiles of pol III on all the tRNA genes; the highest pol III levels at the +40 bp position (61) and two transient pausings at the beginning of the A and B boxes (81), are consistent with the pol III pausing at the genes in the wild type cells. Therefore, similar to earlier reports (36, 77) the observed increase in pol III occupancy (Fig. 7A, 7B and S7A) may be due to the release of pol III pausing in the *paf1*Δ cells. Paf1 has been found to both promote and reduce the pol II pausing (36) on the genes. The recruitment of Paf1 facilitates the Ser2 phosphorylation of pol II, resulting in release of paused pol II into elongation mode. When Paf1 is depleted, the paused pol II levels at 5’-ends increase (78). Pol III is not known to undergo phosphorylation-dependent initiation-elongation mode switching, no factors equivalent to those participating in this switching of pol II are known to affect pol III transcription and pol III-transcribed genes are too short to differentiate between the two modes. Therefore, based on our results we speculate that the PAF1C localization near the TSS in the wild type cells interferes with the progression of the pol III during transcription, resulting in its transient pausing near the A box, seen earlier in the genome-wide studies (61, 81). In the absence of Paf1, high transcription rate and recycling (20, 21) of pol III result in the overloading and obstruction of the short gene body with pol III, eventually providing only a small gain in the transcription. Thus, Paf1 restricts the pol III levels on the tRNA genes by promoting pol III pause at the 5’ gene-ends. While TFIIIC bound to the intragenic boxes A and B may cause the transient pausing of pol III at all the genes (18, 81), the Paf1 presence may increase the pausing near the TSS of the genes in a gene-specific manner.

By promoting the pol III pausing, Paf1 regulates pol III advancement on the genes, which not only reduces pol III enrichment/occupancy and transcription but also allows timely clearance of the short gene body without affecting the high transcription frequency. This avoids the crowding of the short gene length with the transcribing pol III molecules, which may otherwise cause further obstruction to the replication fork progression on these naturally occurring replication fork pause sites. As opposed to this, when pol III is lost from the genes under repression (19, 61), Paf1 is also partially lost. Therefore, the significance of Paf1 association with the pol III TC is to keep pol III levels low on the tRNA genes, which are under replication stress even in the normally cycling cells.

### PAF1C protects tRNA genes against replications stress-associated DNA damage

Pol III TC and transcribed genes are potentially highly active and transcription to their full capacity may be an extravagance for cellular energy. The high transcription activity also makes them vulnerable to replication stress-induced DNA damage (6). High accumulation of pol III on these short genes may also increase the transcription-replication conflict due to the steric clashes between the movements of the DNA and RNA polymerases on the genes. Paf1 intervention by restricting pol III levels on the genes resolves this as well.

The earlier reported DNA pol2 increase on several tRNA genes in *rrm3* cells was found correlated with replication fork pausing (76). Therefore, the increased pol2 levels upon HU exposure and in the *paf1*Δ cells denote slowing down of replication fork at tRNA genes, since the Paf1 deletion is also reported to impair replication fork progression (72). An increase in γ-H2A levels represents replication stress due to the replication slow-down and fork stalling (73). As Paf1 is found to counter the HU-induced replication stress in this study, a reason behind HU sensitivity of the *paf1*Δ cells could be the high increase of pol III levels at the tRNA genes. Reducing the pol III occupancy may be a cellular response to the replication stress caused by the genotoxins in order to reduce the transcription rate and facilitate the replication fork progression on the tRNA genes.

PAF1C cooperates with Mec1 and INO80 in dislodging pol II from the sites of stalled replication fork under replication stress (72). Deletion of the adaptor protein in the replication stress pathway, Mrc1 was found to show an effect similar to Paf1 on pol III occupancy and primary transcript synthesis on some of the tRNA genes (75). However, on the tested pol III-transcribed genes in this study, increased levels of pol III, DNA pol2 and γ-H2A in the absence of PAF1C suggest that in the wild type cells PAF1C normally displaces pol III at the tRNA genes, which are known sites of high transcription activity, replication fork stalling and genomic fragility (6, 7, 89).

The association of proteins and complexes involved in various basic life processes with pol III TC underscores the possibility of the pol III TC serving as a signaling hub for communication between the transcription and other cellular physiological activities. Further studies with many of these newly identified interacting partners may reveal novel cellular stress response pathways, which may link regulation of transcription by all the three RNA polymerases. Our results find little relevance of PAF1C to the known roles of PAF1C in pol II transcription, nucleosome dynamics or histone methylations. The inhibitory influence of PAF1C on the pol III occupancy and replication stress-induced DNA damage on tRNA genes lay emphasis on a protective role for it in the wild type cells. Keeping low pol III levels may reduce the fork stalling, facilitate the fork progression and increase the genome integrity at individual gene loci.

### Data availability

The mass spectrometric proteomics data have been deposited to the ProteomeXchange Consortium via the PRIDE partner repository with the dataset identifier PXD003748.

## Acknowledgements

We thank David Schneider for deletion strains of Paf1 subunits, David Donze for Tfc1-Flag and Kevin Struhl for Tfc6-Flag strains of yeast. Proteomic experiments were partly supported by the Agence Nationale pour la Recherche (ProFi grant ANR-10-INBS-08-01). DVV, AS and PB had been recipients of Senior Research fellowship from UGC, ICMR and CSIR, Govt. of India, respectively.

## Author Contribution

IP samples were prepared by DVV for pol III and by PB for TFIIIC and TFIIIB. Both analysed proteomic data. DVV and PB constructed yeast strains and PB performed Paf1 experiments. AS carried out genome-wide NGS data analyses, BG performed bioinformatics and statistical analyses of the interactome data, YC carried out mass spectrometric runs, data analyses and protein identifications. PB# conceived the study, guided and coordinated the study, compiled the results and wrote the manuscript.

